# Sex-specific behavioral features of juvenile and adult haploinsufficient *Scn2a^+/−^* female mice, model of Autism Spectrum Disorder

**DOI:** 10.1101/2023.12.11.571165

**Authors:** Wendy Marcantonio, Martina Simonti, Isabelle Léna, Massimo Mantegazza

## Abstract

Variants of the *SCN2A* gene, encoding the Na_V_1.2 sodium channel, cause a spectrum of neurodevelopmental and epileptic disorders, and are among those that show the strongest association with Autism Spectrum Disorder (ASD). ASD has a male-bias prevalence, but several studies have proposed that female prevalence may be underestimated due to different symptomatic expression compared with males. However, it is unclear whether this is related to actual different pathological features or to greater masking abilities in females.

Studies on *Scn2a^+/−^* mice, a model of *SCN2A* haploinsufficiency and ASD, have shown an age-dependent ASD-like phenotype attenuated at adulthood in males. However, little is known about the behavioral features of *Scn2a^+/−^* female mice. We performed a battery of behavioral tests that are relevant for assessing ASD-like features, investigating juvenile and adult *Scn2a^+/−^* female mice.

Our results demonstrate that female *Scn2a^+/−^* mice exhibit an overall milder phenotype than males, showing increased sociability and increased risk taking in juveniles, hyper-reactivity to cold stimuli in adults, altered decision-making related behaviors in both. Thus, this is consistent with the male-bias prevalence of ASD and the existence of different ASD phenotypic features in males and females. Both genders should be investigated in studies of mouse models of ASD.

## Introduction

Autism Spectrum Disorder (ASD) is a group of neurodevelopmental dysfunctions with onset in early childhood, characterized by social interaction and communication deficits, restricted and repetitive behaviors, reduced interests, sensory abnormalities and cognitive alterations^1–3^. The refinement of diagnostic criteria and the early assessment of patients have led, over the last few decades, to a significant increase in the number of cases identified, with ASD affecting 1 in 44 children in the United States, across all ethnic and socio-economic groups^4^. Notably, ASD shows a male-oriented imbalance, with a male/female ratio between 3 and 4 in children^3,5,6^. However, it has been shown that women may express ASD symptoms differently than men^7,8^. It has been proposed that this differential manifestation of ASD symptoms in females makes their initial identification more difficult, leading to a delayed or incorrect diagnosis and an underestimation of ASD cases in females^3,9–11^. However, it’s unclear whether this gender difference in ASD phenotype is related to actual differences in pathological features and whether it has actual biological causes. In fact, ASD masking behaviors (mimicry, mirroring of peer behavior, camouflage, etc.) have been shown to be more prominent in females and could attenuate/modulate the symptoms^9,12–15^, which nonetheless may be similar to those of males in their basic features.

Although detailed pathological mechanisms of ASD are not clear yet, both genetic and environmental factors play a role in ASD etiology and it has been proposed that an imbalance between excitation and inhibition in brain circuits may be involved^2,16,17^. Among the genetic causes of ASD, a relevant target is the *SCN2A* gene, encoding the Na_v_1.2 voltage-dependent sodium channel. *SCN2A* genetic variants cause a wide spectrum of neurodevelopmental and epileptic disorders, and they are among the variants that show the strongest association with ASD^18–20^. It has been proposed that *SCN2A* variants causing ASD lead to Na_V_1.2 loss of function (LoF) and haploinsufficiency^18,20,21^. *Scn2a* heterozygous knock-out (*Scn2a^+/−^*) mice^22^, which model *SCN2A* haploinsufficiency, show an age-dependent ASD phenotype attenuated in adulthood, associated with mild absence-like seizures and memory impairments^23–26^. Some of the phenotypic features observed in *Scn2a^+/−^* mice are also present in other mouse models of *Scn2a* haploinsufficiency and autism^27–29^. Interestingly, increased sociability and reduced anxiety-like behavior are the most recurrent behavioral features observed in *Scn2a* mouse models^26,28–30^. Additionally, different types of stereotyped repetitive behaviors have been reported^23,26,28,31^.

Historically, behavioral studies were performed using mainly male rodents to avoid interferences with estrus cycle and female hormonal modulations occurring during the tests. Also with regard to *Scn2a* mouse models, only a few studies have evaluated some behavioral features of females, without carrying out extensive characterizations^27–29,31^. Here, we performed on female *Scn2a^+/−^* mice a battery of behavioral tests relevant to the assessment of ASD-like features. To evaluate the developmental characteristics of the phenotype, we studied *Scn2a^+/−^* and wild type (WT) littermate female mice in the juvenile (post-natal day, P, P21-44) and the adult (>P60) period, and compared these results with those that we already obtained with *Scn2a^+/−^* male mice^23^.

## Material and methods

### Animals

The generation and the genotyping of *Scn2a^+/−^* mice have been described by Planells-Cases et al.^22^. The original heterozygous breeders were provided by Dr. E. Glasscok of Louisiana State University Health Sciences Center (Shreveport). Heterozygous (HTZ) *Scn2a^+/−^* and control *Scn2a^+/+^* littermate mice were generated from crosses between HTZ males and wild-type (WT) females on a C57Bl6/J background (Charles River, France). All mice were housed in a controlled environment (22–24°C, 40– 50% humidity) under a 12h dark/light cycle with access to food and water ad libitum. After weaning (post-natal days (P) 21–23), mice were group-housed with same sex littermates of mixed genotype (n=3–5/cage) in standard polycarbonate cages containing plastic houses, wood sticks and nesting material. In this study we performed an extensive behavioral characterization of female mice and selected tests on male mice. Behavioral tests were performed sequentially in juvenile (between P21 and P44, corresponding to the pre-pubescent, P21–34, and pubescent, P35–44, periods of adolescence ^32,33^ and adult (between P60 and P95) female mice. For all experiments, the animals were habituated to the testing room at least 30 min before the beginning of the tests. The animals were submitted to the behavioral tasks in the following order: Elevated Plus Maze (EPM), openfield, 3-chamber social interaction, Y-maze, grooming, olfaction, marble burying, novel object recognition, rotarod, hot and cold plate. All the tests were conducted on the same mice, from juvenile to adulthood stages, except the hot and cold plate tests that have been performed on distinct groups of mice to avoid cross influence of potential skin damages induced by the first plate tests. A total of 5 litters have been used to assess the behavioral phenotype of female *Scn2a^+/−^* mice (for a total of 21 WT and 23 HTZ mice) and 2 litters for testing male *Scn2a^+/−^*mice (for a total of 27 WT and 28 HTZ). Mice were randomized (https://www.randomizer.org/); experiments and analysis were performed blind to genotype by two experimenters (W.M. and M.S.). All procedures were performed in accordance with the European Council Directive (2010/63/EU) and approved by the local ethics committee (Comité Institutionnel d‘Éthique Pour l’Animal de Laboratoire (CIEPAL)-Azur, https://ciepal-azur.unice.fr/) and the French Ministry of Research (http://www.enseignementsup-recherche.gouv.fr/, APAFIS#2023020813272228).

### Open field test

The subject mouse was allowed to explore a white wooden box (40×40×40cm) for 30 min under anxiogenic lighting (300 Lux). The apparatus is virtually divided in 3 zones: periphery (5 cm to the edges), center (20×20cm) and intermediate zone. Distance travelled and time spent in periphery and center zones was assessed by automatic tracking the animals with the ANY-maze software (Stoelting, Europe, Dublin, Ireland). To evaluate anxiety-like behavior, percent of time spent in the center versus the periphery was calculated. Entries into a zone were automatically measured, considering an entry when 90% of the body was in the zone and an exit when at least 50% was out.

### 3-chamber sociability test

The 3-chamber test is used to assess sociability and social memory in mice. It was performed in both juvenile (P23-28) and adult (>P70) females under low light (30 Lux). The apparatus is a grey rectangle plastic box divided in three zones of equal size (30×35×35cm) separated by transparent walls with 2 openings. The two lateral chambers contain a pierced (ø10mm holes) transparent cylinder (8cm diameter) which allows physical contacts between mice. This test consists of 3 phases of 10 minutes: Habituation, Sociability and Social Memory^34,35^. Briefly, the subject mouse was initially allowed to freely explore the apparatus for 10 min during the habituation phase. Then, it was placed in the center zone while a stranger mouse (named “unfamiliar”, S1) was placed in one of the two empty cylinders (in the side less preferred in the habituation phase) and allowed to explore again for 10min. In the third phase, the tested mouse was placed again in the center zone and a second new unfamiliar mouse was added in the other cylinder (named “unfamiliar”, S2, whereas the previous one is considered in this phase as “familiar”); the tested mouse was then allowed to explore for 10 more minutes. Tracking of movements and assessment of the time spent in each chamber was done with ANY-maze. Close interactions with empty cylinders, unfamiliar or familiar mice (time spent sniffing closely) were manually assessed and the total time in close interaction calculated (Unfamiliar + Empty, T_S1+E_ or Familiar + Unfamiliar, T_S1+S2_). Social Discrimination Index (SDI) and Social Memory Index (SMI) were calculated as: SDI= (T_S1_-T_E_)/T_S1+E_; SMI=(T_S2_-T_S1_)/T_S1+S2_. A positive index is a sign of social preference or valid social memory, as the tested mouse spent more time with the social or unfamiliar stimulus, respectively. Negative index indicates aversive behavior towards the social stimulus. A score of 0 indicates no preference or no memory of the previously encountered stimulus mouse. Entry into a zone was automatically calculated if 90% of the body was in the zone and an exit if at least 50% was outside.

### Olfactory discrimination

Olfactory discrimination between social and non-social odors, which could interfere with the social behaviors measured in mice, was assessed as previously described with minor modifications^23,36^. Squares of filter paper (5×5 cm) were dampened with water (neutral control) or coffee (non-social aversive odor) (approximately 0.5ml). Social odors (attractive) were obtained placing filter paper squares for 3 days in cages housing non familiar juvenile or adult females. Mice were placed in clean plexiglas cages without bedding for 10 min of habituation. Then, filter papers were introduced sequentially (water, coffee, social odor) for 3min each, spaced by 1min interval. Olfactory preference was assessed as the difference between the time spent interacting with (sniffing) social or non-social odors and that spent interacting with (sniffing) the paper dampened with water.

### Rotarod

Motor coordination, balance and motor learning were evaluated using an accelerating rotarod (Bioseb-*In vivo* Research Instruments, Vitrolles, France)^37^. Mice were habituated to the procedure before the test phase: each mouse was trained until able to stay 1min on the circular rod at a constant speed of 4rpm. Mice were then placed on the rod that accelerated from 4 to 40 rpm over 5min. Mice were tested with three trials a day (inter-trial interval of 30 min) for 2 consecutive days. Fall latency from the rotating rod was automatically recorded; mice that fell before 15 seconds were put back on the rod and retested.

### Marble burying test

We used the marble burying (MB) test to evaluate motor stereotypies and repetitive behaviors^38^. The apparatus consists of a transparent plastic arena (30×30×30cm) filled with 5 cm of clean bedding (poplar wood chips, average particle size 3.5–4.5 mm; Lignocel Select, SAFE, France) on which 20 marbles are arranged (5×4 rows). The tested mouse was allowed to explore the apparatus for 20 min and the number of marbles buried every 5 min was manually scored by the experimenter. A marble was considered buried if more than 50% was buried in the bedding, not counting again the same marble if buried several times.

### Grooming

We performed the self-grooming test in an open field (40×40×40 cm) arena to further study repetitive behaviors. The mouse is placed in the open field in low lighting (30 Lux – non anxiogenic) for 20 min. After 10 min of habituation, the time spent spontaneously self-grooming was measured manually.

### Elevated Plus Maze

We used the Elevated Plus Maze (EPM) test to investigate anxiety-like behavior. The apparatus consists of a grey plastic “plus-shaped” cross (25×5×2.5 cm) elevated 1 meter above the ground, with 2 open arms (anxiogenic zone) and 2 enclosed arms (non-anxiogenic zone), separated by a small center zone (neutral zone). The tested mouse was allowed to freely explore the maze for 5 min. Normally mice spend more time in non-anxiogenic zones than in the anxiogenic zones, and modifications of anxiety levels can modulate this parameter. We scored automatically with the ANY-maze tracking software distance, speed, time spent and number of entries in each zone, considering 90% and 50% of the body for entry and exit parameters, respectively.

### Hot and Cold Plate tests

Thermal sensory processing was evaluated by the hot and cold static plate tests in distinct cohorts. The experimental protocol was as in (Corder et al. 2017; Eaton et al. 2021), with little modifications. Briefly, the tested mouse was placed on a 30°c plate (Bioseb-*In vivo* Research Instruments, Vitrolles, France) for a 5 min habituation phase. After 30 min, the mouse was placed back on the plate, which was heated to 52°C or cooled to 7.5 °C, the latency to the first sign of discomfort was manually scored and the mouse removed from the plate. Signs of discomfort taken into account were hind paw lifting off the plate (>2 sec), shaking, licking, or jumping on the plate. If no sign of discomfort was detected after a cut-off time of 45s (hot plate) or 1 min (cold plate), the mouse was removed from the device to avoid skin injuries. The analysis was performed on video recordings from the top.

### Novel Object Recognition Test (NORT)

This memory test based on the spontaneous tendency of rodents to explore novelty has been performed as previously described^39^ in an open-field apparatus slightly illuminated (30 lux). During the phase of habituation (Day 1), mice were allowed to freely explore the arena for 10min. On the experimental day (Day 2), mice were submitted to two trials with an inter-trial interval of 1hour. During the first trial (acquisition phase), the tested mouse was placed in the open-field in the presence of two identical objects (wavy transparent glass square saltshakers) and allowed to freely explore for 10min. During the second trial (retention phase), one of the objects was replaced by a novel object (opaque smooth black round ceramic cup) and the mouse was allowed to explore both objects (novel and familiar) for 5min. Mice that did not explore the objects for a minimum of 15s were excluded (juveniles: 3WT and 1 HTZ; adults: 3WT and 4HTZ). The pairs of objects used were different in shape, color and texture, but had similar sizes. The two objects were placed symmetrically in the center of the arena about 6 cm away from the wall. The location of the novel object was counterbalanced across animals (right or left side). Recognition memory was evaluated using a discrimination index (DI), quantified as the difference between the time spent exploring the novel object and the familiar object, divided by the total time of exploration of both objects and expressed as percentage.

### Y maze test

Spatial working memory was assessed quantifying the spontaneous alterations in a symmetrical grey plastic Y maze (30×6×15cm), in which the tested mouse was placed at the end of one arm and allowed to freely explore the maze for 8min^40,41^. The sequence and the number of arm entries were automatically quantified with the ANY-maze software, considering 99% of the body for entry and 50% for exit. An alternation was defined as consecutive entries into the three different arms (A, B or C) on overlapping triplet sets (ex: ABCABBACAB=5 alternations). The spontaneous alternation score (%) was calculated as the ratio of actual (total alternations) to possible (total arm entries-2) alternations, expressed as percentage. The percentage of time spent in the center zone while moving between two arms (considered as a decision-making parameter) and the total distance travelled were also automatically evaluated.

### Statistical analysis

Data were analyzed for normality (Shapiro-Wilk test) and equal variances using GraphPad Prism 10.0.3 (GraphPad Software, San Diego, California USA). When data were normally distributed, comparisons between genotypes in young or adult mice were evaluated using unpaired Student’s t-test or Welch’s t-test (when variances were different); the Mann-Whitney test was used for non-normally distributed data. When appropriate, a two-way (repeated) measures ANOVA followed by Sidak’s multiple comparison post-hoc test was used. One sample Wilcoxon t-tests were used to compare the discrimination index with a 0% chance level in the novel object recognition test and the 3-chamber sociability test. The results are displayed as mean±S.E.M. We considered the differences as statistically significant when p≤0.05, * indicates a significant difference between genotypes and # indicates a significant difference in comparisons within subjects (olfaction test) or with chance level. We considered borderline significance (strong trend) when p-values were between 0.05 and 0.09. plots were generated with GraphPad Prism 10.0.3 or Excel.

## Results

### Female *Scn2a^+/−^* mice show normal locomotor activity and no anxiety-like behavior in the openfield

We initially assessed locomotor activity because it is an important feature that can influence the outcome of numerous behavioral tests. We quantified it in the highly illuminated arena of the open field apparatus during 30min-long sessions, considering the whole arena, the periphery or the center (Figure 1A). The locomotor activity showed no difference between juvenile female *Scn2a^+/−^* (HTZ) mice and WT female littermates (Figure 1B), in the whole arena (66.9±2.4m distance traveled for HTZ, 67.3±5.1m for WT; Mann-Whitney test: U=201, p=0.3509, NS), the periphery (30.3±2.7m HTZ; 31.2±1.2m WT; Mann-Whitney test U= 183, p=0.17, NS) or the center (11.5±1.1m HTZ; 11.1±0.8m WT; Student’s t-test Center: t(42)=0,3447, p= 0,73, NS). We obtained similar results for adult females (Figure 1D) in the whole arena (58.4±3.9m HTZ, 63.4±4.7m WT, Student’s t test: t(36)=0,8240, p=0.41, NS), the periphery (29.7±1.8m HTZ; 32.3±2.6m WT, Student’s t test: t(36)=0,8683, p=0.40, NS) or the center (7.3±0.7m HTZ; 8.0±0.7m WT; Student’s t test, t(36)=0.6330, p=0.53, NS). Moreover, we evaluated the anxiety behavior by analyzing the percentage of time spent in the center zone of the open field arena (Figure 1C-E), which showed no difference, neither for juvenile females (13.9m3±1.9% HTZ, 12.9m1±1.6% WT; Mann-Whitney test: U=239.5, p=0,97, NS) nor for adult females (6.3±1.0% HTZ, 7.4±0.8% WT, Student’s t test: t(36)=0,8176, p=0.4190, NS). Therefore, *Scn2a^+/−^*female mice show normal exploratory and locomotor activity and no anxiety-like behavior across development.

**Figure 1:**
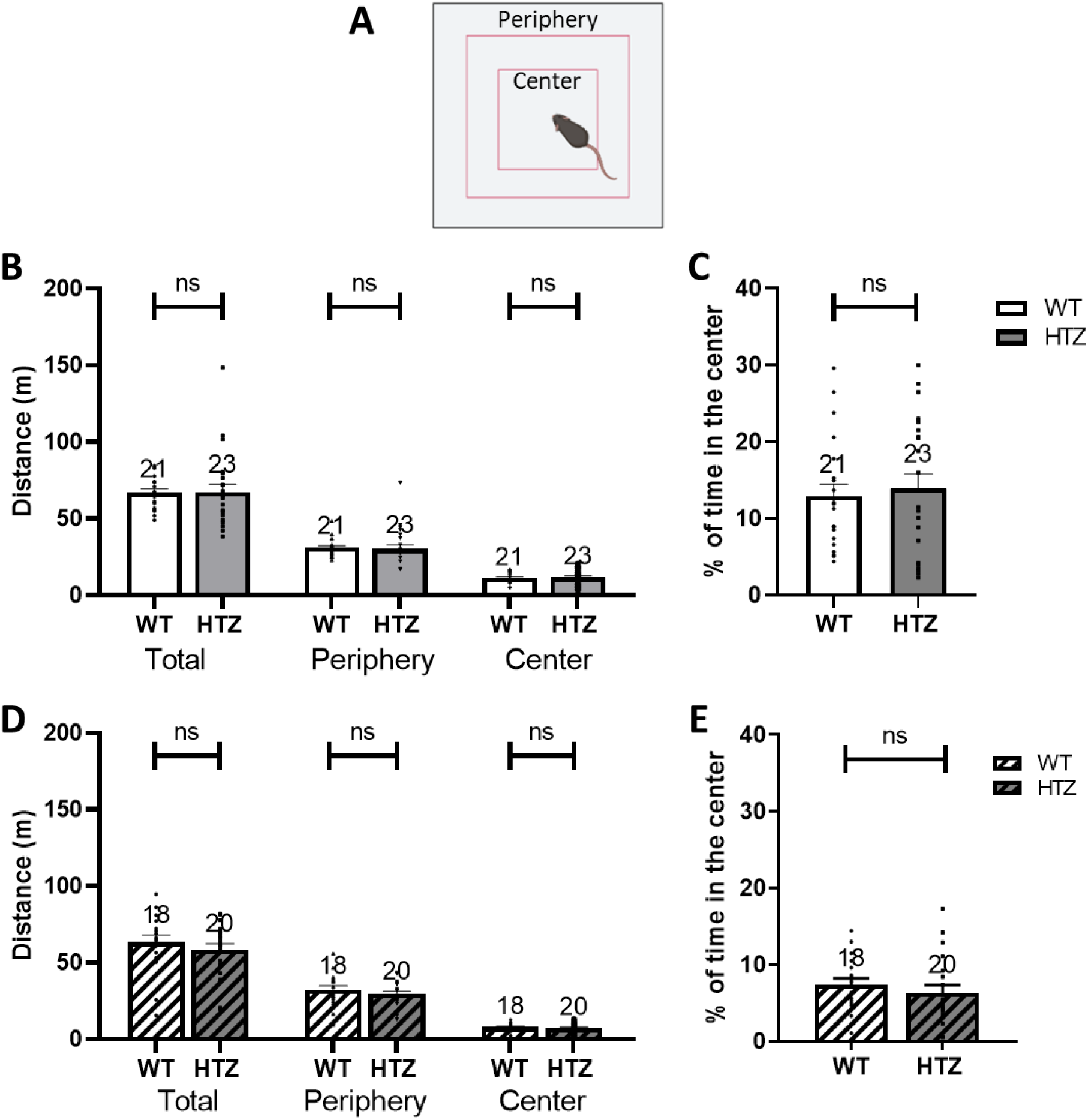
Evaluation of locomotor activity in juvenile and adult *Scn2a*^+/−^ (HTZ) female mice. **A:** Schematic illustration of the apparatus, which for the analysis was virtually divided in zones: center, intermediary and periphery. **B-C:** Locomotor activity in 30min (total, periphery and center) (B) and percent time spent in the center of the open field in 30min (C) for juvenile female mice. **D-E:** Locomotor activity in 30min (total, periphery and center) (D) and percent time spent in the center of the open field in 30min (E) for adult female mice. ns=non-significant (Mann-Whitney or Student’s t-test); the number of mice (N) is indicated above the bars. White: WT, Grey: HTZ. No Pattern, juvenile; striped, adult.

### *Scn2a^+/−^* female mice show abnormal sociability but no alteration of social memory in the 3-chamber test

Abnormality in social interactions is a major clinical feature in ASD. The 3-chamber test (three connected chambers, in which transparent holed cylinders are placed in the two opposite side chambers) is used to evaluate sociability in rodents, exploiting their natural preference for social stimuli and social novelty. Sociability and social memory are assessed by the time spent in close interaction with the two cylinders (empty vs. unfamiliar mouse or familiar mouse vs. unfamiliar mouse, respectively) (Figure 2A-B). Sociability and social memory indexes are calculated by subtracting the time spent in interaction with the empty cylinder or the familiar mouse cylinder, respectively, to the time spent in interaction with the unfamiliar mouse cylinder, and by dividing it by the total time spent in close interaction with both cylinders. We tested juvenile and adult females and settled as exclusion criterion a minimum of 40-second of total interaction with both cylinders. Two WT and one HTZ juvenile mice and one WT adult mouse were excluded from the analysis because of this criterion.

**Figure 2:**
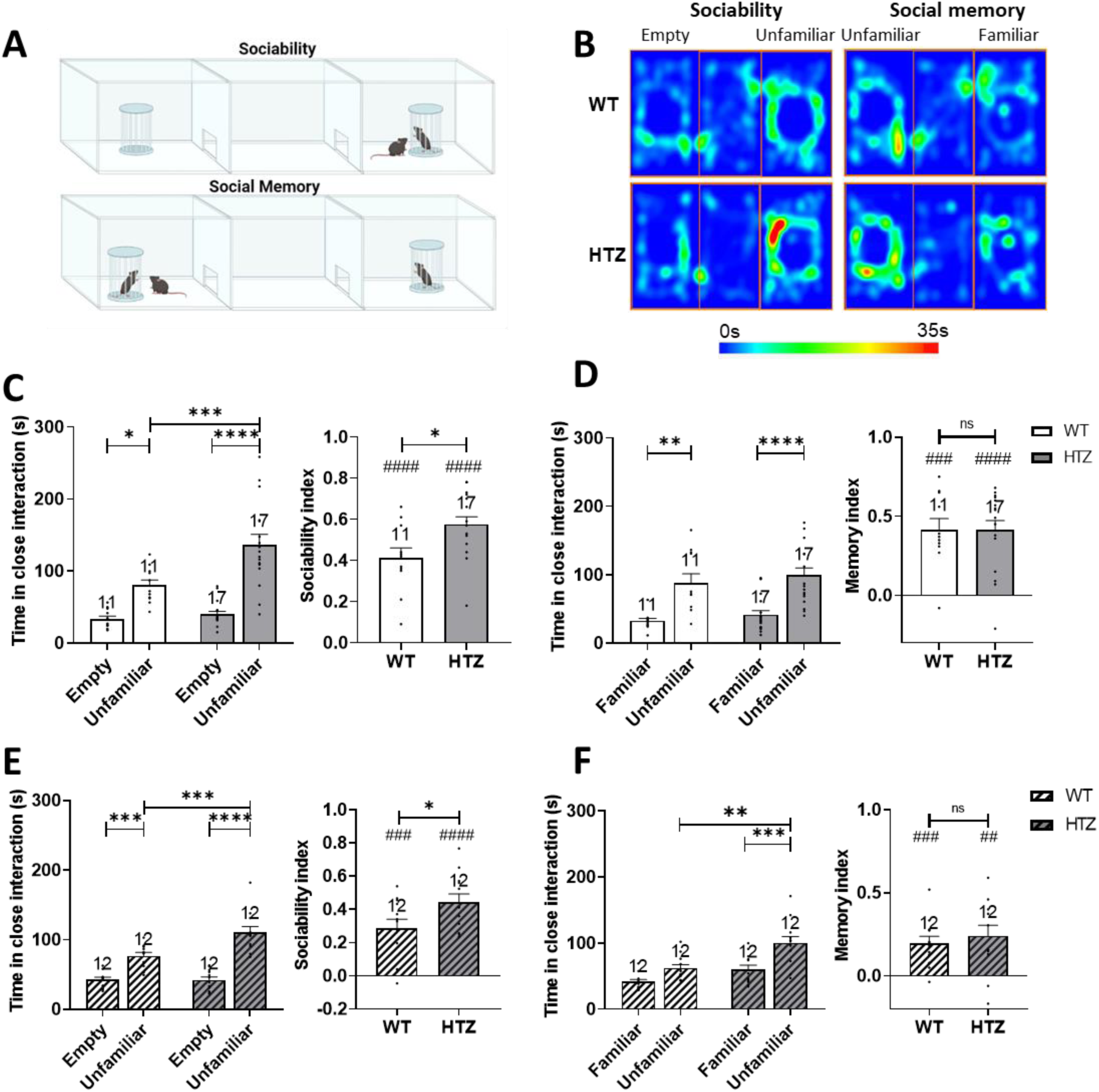
Evaluation of sociability in juvenile and adult *Scn2a*^+/−^ (HTZ) female mice. **A:** Schematic diagram of the 3-Chamber Sociability and Social Memory tests. **B:** Representative heatmap of the time spent by WT and HTZ mice in different areas of the apparatus during the Sociability and Social Memory phases. The red areas indicate the locations where the test mouse spent the maximum amount of time. **C-D:** Results in juvenile HTZ female mice for the sociability and social memory phases of the test: (C) Time in close interaction with the empty cylinder or the unfamiliar mouse (Left) and sociability index (Right) in juvenile female mice. (D) Time in close interaction with the familiar or the new unfamiliar mouse (Left) and social memory index (Right) in juvenile female mice. **E-F:** Results for adult HTZ female mice for sociability or social memory phases of the test. (E) Time in close interaction with the empty cylinder or the unfamiliar mouse (Left) and sociability index (Right) in adult female mice. (F) Time in close interaction with the familiar or the new unfamiliar mouse (Left) and social memory index (Right) in adult female mice. Statistical analysis: Two-way ANOVA with Sidak’s post-hoc test for multiple comparison test, Mann-Whitney or Student’s t-test, *p<0.05, **p<0.01, ***p<0.001, ****p<0.0001. One-sample t-test for comparison to chance level #p<0.05, ##p<0.01, ###p<0.001, ####p<0.0001. The number of mice (N) is indicated above the bars. White: WT, Grey: HTZ. No Pattern, juvenile; striped, adult.

For juvenile females, we observed a significant increase in the time spent in close interaction with the stimulus mouse compared to the empty cylinder for both genotypes in the sociability phase, but the increase was higher for HTZ mice (33.3±4.1s WT empty, 80.0±7.5s WT with unfamiliar, 39.6±4.2s HTZ with empty, 136.5±14.2s HTZ with unfamiliar; two-way ANOVA, F (1, 52) = 53,20, P<0,0001 for stimulus effect, F(1.52)=10.16, p=0.0024 for genotype effect and F(1.52)=6.478, p=0.0139 for genotype x stimulus interaction; Sidak’s post-hoc test: WT empty vs unfamiliar p=0.018, HTZ empty vs unfamiliar p<0.0001; WT unfamiliar vs HTZ unfamiliar p=0.0009) (Figure 2C, left). Consistently, the sociability index was significantly higher than chance (index = 0) level for both genotypes (0.58±0.04 HTZ 0.41±0.05 WT One-sample t-test, t(10)=8.132, p<0.0001, for WT, t(15)=14.79, p<0.0001 for HTZ), but higher for HTZ females compared to controls (Student’s t test, t(25)=2.527, p=0.018) (Figure 2C, right). In the social memory phase, both HTZ and WT juvenile female mice exhibited a social preference for the unfamiliar mouse (cylinder effect) and, differently than for the sociability phase, there was no significant difference between genotypes in the time spent in close interaction with the unfamiliar mouse (98.9±10.9s HTZ with unfamiliar; 41.1±6.1s HTZ with familiar; 87.6±13.7s WT unfamiliar; 32.2±3.8s WT familiar; two-way ANOVA, F(1.52)=34.54, p<0.0001 for cylinder effect, F(1.52)=1.107, p=0.2976 for genotype effect, NS and F(1.52)=0.01615, p=0.9 for genotype x cylinder interaction) (Figure 2D, left). Consistently, the social memory index was different from to chance level for both groups (0.42±0.06 HTZ; 0.42±0.07 WT; one-sample t-test: t(10)=5.918, p=0.0001 for WT compared to chance level, t(16)=6.779, p<0.0001 for HTZ compared to chance level), but not significantly different between HTZ and controls (Mann-Whitney, U=88.50, p=0.8262) (Figure 2D, right).

The results that we obtained for the sociability phase of adult females are similar to those of juvenile females. We observed a significant increase in the time spent in close interaction with the stimulus mouse compared to the empty cylinder for both genotypes, but the increase was higher for HTZ mice (42.2±4.3s HTZ with empty; 110.8±8.4s HTZ with unfamiliar; 42.5±3.7s WT with empty; 76.8±4.8s WT with unfamiliar; two-way ANOVA, F(1.44)=84.78, p<0.0001 for stimulus effect, F(1.44)=9.031, p=0.0044 for genotype effect and F(1.44)=9.420, p=0.0037 for genotype x stimulus interaction; Sidak’s post-hoc test: WT empty vs unfamiliar p=0.0005, HTZ empty vs unfamiliar p<0.0001 WT unfamiliar vs HTZ unfamiliar p=0.0006) (Figure 2E, left). The sociability index was significantly higher than chance level for both genotypes (0.44±0.05 HTZ; 0.29±0.06 WT; one-sample t-test t(11)=5.163, p<0.0003 for WT, t(11)=8.939, p<0.0001 for HTZ), but higher for HTZ females compared to controls (Student’s t test: t(22)=2,115, p=0.0460) (Figure 2E, right). The multiple comparison of the time spent in close interaction during the social memory phase showed a significant preference for the new unfamiliar mouse compared to the familiar one in HTZ adult females, whereas WT females showed a trend toward preference for the new unfamiliar mouse (100.1±10.0s HTZ with unfamiliar; 60.1±6.6s HTZ with familiar; 62.3±5.2s WT with unfamiliar; 41.5±3.5s WT with familiar; two-way ANOVA, F(1.44)=20.19, p<0.0001 for cylinder effect, F(1.44)=17.47, p=0.0001 for genotype effect and F(1.44)=2.016, p=0.1627 for genotype x cylinder interaction; Sidak’s post-hoc test: WT familiar vs unfamiliar p=0.19, HTZ familiar vs unfamiliar p=0.0008, WT unfamiliar vs HTZ unfamiliar p=0.0016) (Figure 2F, left). Notably, the social memory index was significantly different than the chance level for both groups (0.24±0.06 HTZ; 0.20±0.04 WT; one-sample t-test t(11)=4.651, p=0.0007 for WT, t(11)=3.781, p=0.003 for HTZ) and there was no significant difference between the index of WT and HTZ females (Student’s t-test, t(22)=0.5906, p=0.56) (Figure 2E, right). Consistent with the trend observed in the multiple comparison test (Figure 2F, left), the comparison of the interaction time of WT mice (familiar vs new unfamiliar mouse) using the paired Student’s t-test showed a significant preference for the new unfamiliar mouse, as expected for WT mice (t(22)=3.288, p=0.0034).

Therefore, both juvenile and adult female *Scn2a^+/−^* mice show hyper-sociability, but no alteration in social memory.

### Olfactory discrimination is functional in *Scn2a^+/−^* female mice at both ages

We assessed olfactory preference and social odor discrimination of *Scn2a^+/−^* female mice, which may be involved in the differences that we observed in sociability. Notably, Na_V_1.2 is expressed also in olfactory granule cells and it has been proposed that it could drive rapid-odor discrimination^42^. Thus, we tested the ability of females to discriminate aversive (coffee) and attractive (social) odors, using the same protocol that we previously used for *Scn2a^+/−^* male mice (quantified as the difference between the time mice spent interacting with social or non-social odor cues and that spent interacting with paper dampened with water)^23^ (Figure 3A).

**Figure 3:**
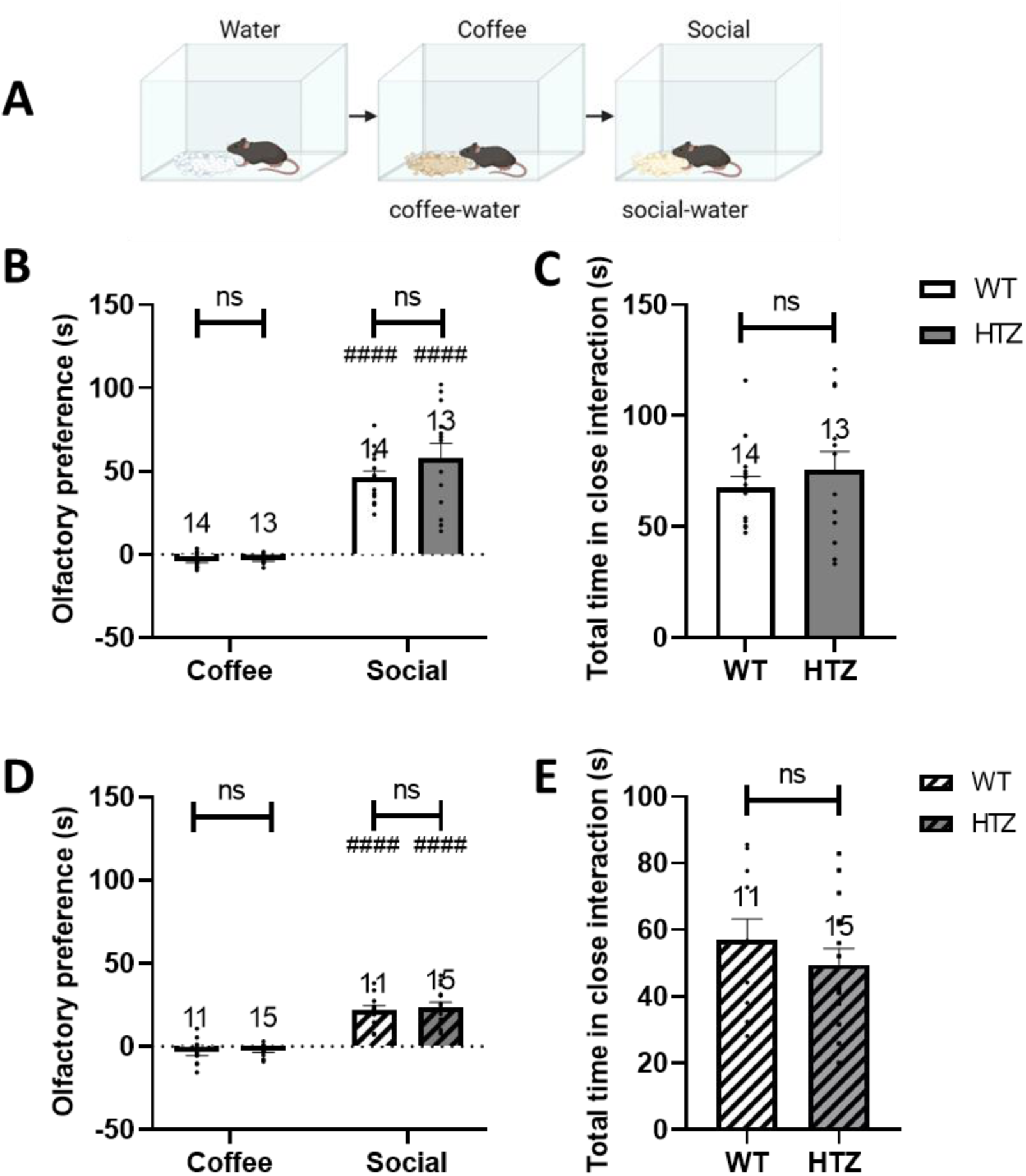
Evaluation of olfactory preference in juvenile and adult *Scn2a*^+/−^ (HTZ) female mice. **A:** Schematic illustration of the olfactory test divided in 3 phases: exploration of paper dampened in water, coffee (aversive odor) and social odors (appetitive odor). Scores are calculated as time spent exploring coffee or social odors minus the time spent exploring water. **B-C:** Olfactory preference in juvenile female mice and total time spent in interaction. **D-E**: Olfactory preference in adult female mice and total time spent in interaction. Statistical analysis: 2 Way Anova, significant odor effect ****p<0.0001 (B-D), Mann-Whitney test: ns (C) and Student’s t-test: ns (E). Post-hoc analysis: Sidak’s multiple comparison test, #### p<0.0001, comparison between coffee and social odors, comparison between genotype: ns. The number of mice (N) is indicated above the bars. White: WT, Grey: HTZ, No pattern: Juvenile, Striped: Adult.

The analysis of the data revealed a significant difference in exploration duration of social vs coffee cues but no inter-genotype effect (two-way ANOVA, juvenile: 58.24±8.6s HTZ social; −3.12±0.8s HTZ coffee; 46.24±1.0s WT social; −3.90±1.0s WT coffee; F(1.50)=141.6, <0.0001 for odor effect, F(1.50)=1.845, p=0.1805 for genotype effect, NS and F(1.50)=1.438, p=0.2361 genotype x odor interaction, NS, Figure 3B; adult: 23.6±3.0s HTZ social; −2.8±0.9s HTZ coffee; 21.7±2.9s WT social; −3.2±2.3s WT coffee; F(1.48)=114.4, p<0.0001 for odor effect, F(1.48)=0.2223, p=0.64 for genotype effect and F(1.48)=0.1032, p=0.75 for genotype x odor interaction, Figure 3D). Post-hoc analysis demonstrated that the comparison between coffee and social cues are significantly different for genotype at both ages (Sidak’s multiple comparison, juvenile (both genotypes): p<0.0001 and adult (both genotypes): p<0.0001, shown as # in the figure). The total time exploring all cues (water + coffee + social) was used as a marker of interest and exploration and showed no statistical difference between genotypes at both ages (juvenile: 75.6±8.4s HTZ; 67.6±5.1s WT; Mann-Whitney, U=79, p=0.58, Figure 3C; adult: 49.6±4.9s HTZ; 57.0±6.2s WT; Student’s t-test, t(24)=0.9536, p=0.35, Figure 3E).

Thus, we observed that both juvenile and adult female mice of both genotypes are able to perform olfactory discrimination equally well. These results show that olfactory discrimination is functional in *Scn2a^+/−^* female mice and therefore is not involved in the altered sociability.

### *Scn2a^+/−^* females do not show clear motor stereotypies, although juvenile females perform better on an acquired Rotarod task

Motor stereotypies associated with restricted interests are among the main diagnostic characteristics in ASD. To investigate these features in *Scn2a^+/−^* female mice, we quantified innate (self-grooming, marble burying) and acquired (Rotarod) motor behaviors (Figure 4A).

**Figure 4:**
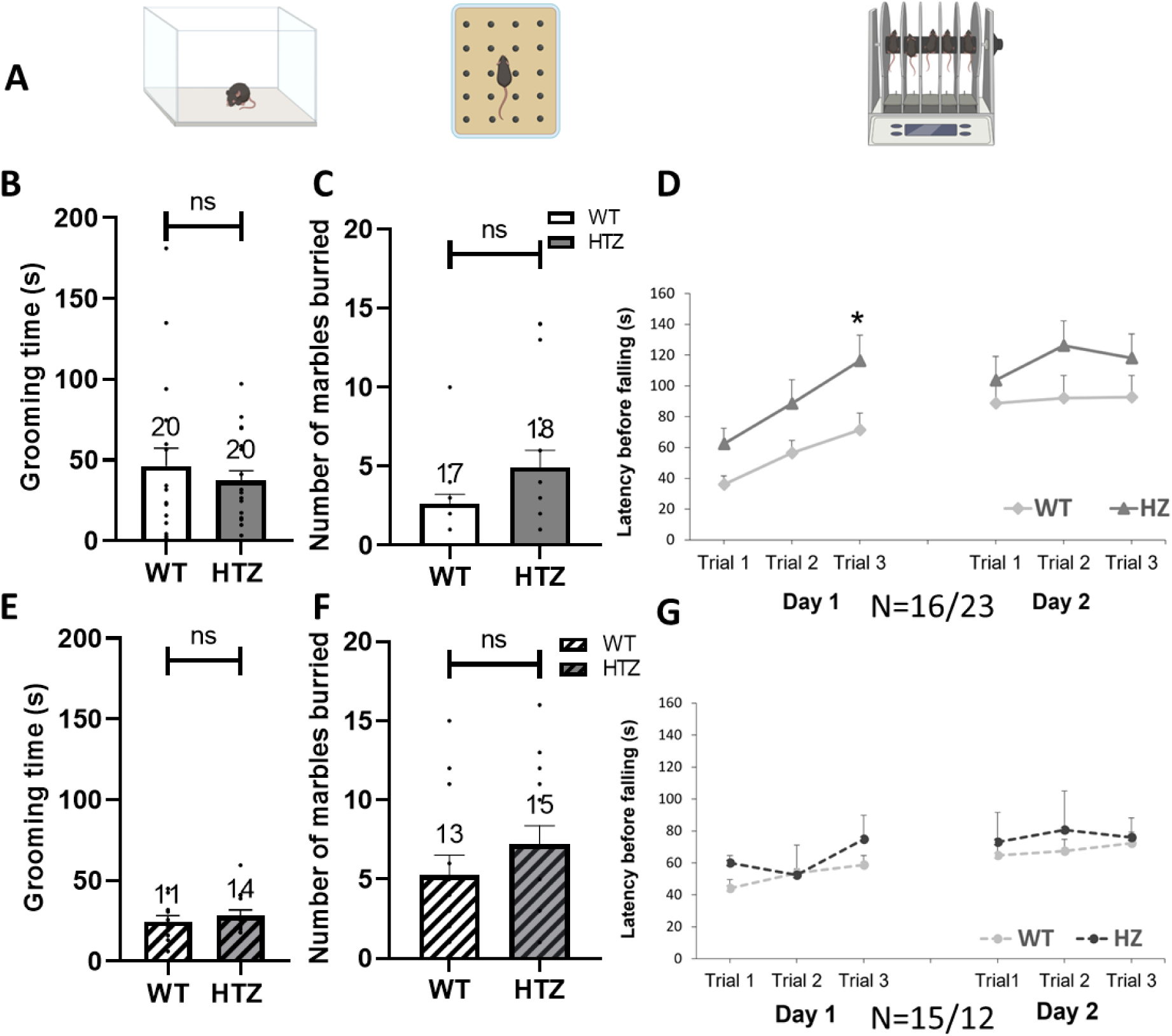
Evaluation of motor stereotypies in juvenile and adult *Scn2a*^+/−^ (HTZ) female mice. **A:** Schematic illustrations of grooming test, marble burying and rotarod apparatuses. **B-D:** Evaluation of innate and acquired motor stereotypies in juvenile female HTZ mice. (B) Time spent self-grooming. (C) Number of marbles buried. (D) Latency to fall from the rotarod, 3 trials per day on 2 days. **E-G:** Evaluation of innate and acquired motor stereotypies in adult female HTZ mice. (E) Time spent grooming. (F) Number of marbles buried. (G) Latency to fall from the rotarod, 3 trials per day on 2 days. Statistical analysis: Mann-Whitney or t-test for grooming and marble burying tests and 2 Way Repeated Measure Anova for rotarod analysis *p<0.05 compared to WT, ns: non-significant. In D the difference is significant considering the two days separately. The number of mice (N) is indicated above the bars or below the plots. White: WT, Grey: HTZ, No Pattern: Juvenile, Striped: Adult.

Neither juvenile nor adult HTZ females showed alterations in the amount of time spent self-grooming (juvenile: 37.3±6.1s HTZ; 46.1±11.0s WT; Mann-Whitney, U=200, p>0.9999, NS, Figure 4B; adult: 28.2±3.7s HTZ; 24.8±3.6s WT; Student’s t-test: t(23)=0.6509, p=0.52, NS, Figure 4E) or in the number of marbles buried (juvenile: 4.9±.1.1 HTZ; 2.6±0.6 WT; Mann-Whitney, U=107.5, p=0,1330, NS, Figure 4C; adult: 7.2±1.2 HTZ; 5.2±1.3 WT; Student’s t-test: t(26)=1.127, p=0.27, NS, Figure 4F). These results indicate that *Scn2a^+/−^* female mice do not display deficits in these innate motor behaviors.

A more pronounced amelioration in performance and faster motor learning in the rotarod test compared to control mice are features considered as stereotypic motor routines in ASD mouse models^43^. To evaluate the motor behavior acquired during the rotarod test, we measured the latency before falling off the rotating and accelerating rod of the apparatus in two daily three-trial sessions. We initially compared the two genotypes including all the trials, and we observed a strong trend towards increased latency in juvenile *Scn2a^+/−^* female mice (two-way repeated measure ANOVA on both days, F(4.177, 154.5)=8.796, p<0.0001 for trial effect, F(1.37)=3.895, p=0.056 for genotype effect and F(5.185)=0.4196, p=0.83 for genotype x trial interaction), but no difference in adult mice (two-way repeated measure ANOVA on both days, F(3.460, 86.51)=4.052, p=0.0068 for trial effect, F(1.25)=0.9008, p=0.35 for genotype effect and F(5.125)=0.51, p=0.7693 for genotype trial interaction). Thus, we performed comparisons for juvenile mice by considering the two days separately. We found a significant genotype effect for day 1 (two-way repeated measure ANOVA on Day 1, F(1.760, 65.12)=10.31 p=0.0002 for trial effect, F(1.37)=5.378, p=0.026 for genotype effect and F(2.74)=0.4599, p=0.6331 for genotype x trial interaction, Figure 4D) but not for day 2 (two-way repeated measure ANOVA on Day 2, F(1.982, 73.32)=0.7268, p=0.48, NS for trial effect, F(1.37)=1.860, p=0.1809, NS for genotype effect and F(2.74)=0.3749, p=0.69, NS for genotype x trial interaction, Figure 4G). These results show that juvenile *Scn2a^+/−^* female mice can perform better on the rotarod than their WT littermates before training, although after training the difference was attenuated. Thus, there were no clear motor stereotypies.

### Sensitivity to cold, but not hot, stimuli is increased in both juvenile and adult male and in adult female *Scn2a^+/−^* mice

ASD patients often show hyper- or hypo-reactivity to sensory (thermal, auditory, visual or tactile) stimuli. Moreover, a recent work has shown that a gene-trap mouse model bearing ∼75% reduction of *Scn2a* expression has an abnormal reactivity to extreme temperatures^27^. We did not investigate these features in our previous study on male *Scn2a^+/−^* mice^23^. Therefore, we evaluated here the reactivity to thermal stimuli in juvenile and adult *Scn2a^+/−^* mice of both sexes, performing the hot (52°c) and cold (7.5°c) plate tests (Figure5A & F). We considered withdrawal (shaking, licking, or lifting) of the hind paws or escaping as markers of thermal discomfort^44^, and selected a cut-off time of 45s for the hot plate and 60s for the cold plate tests to avoid skin lesions.

**Figure 5:**
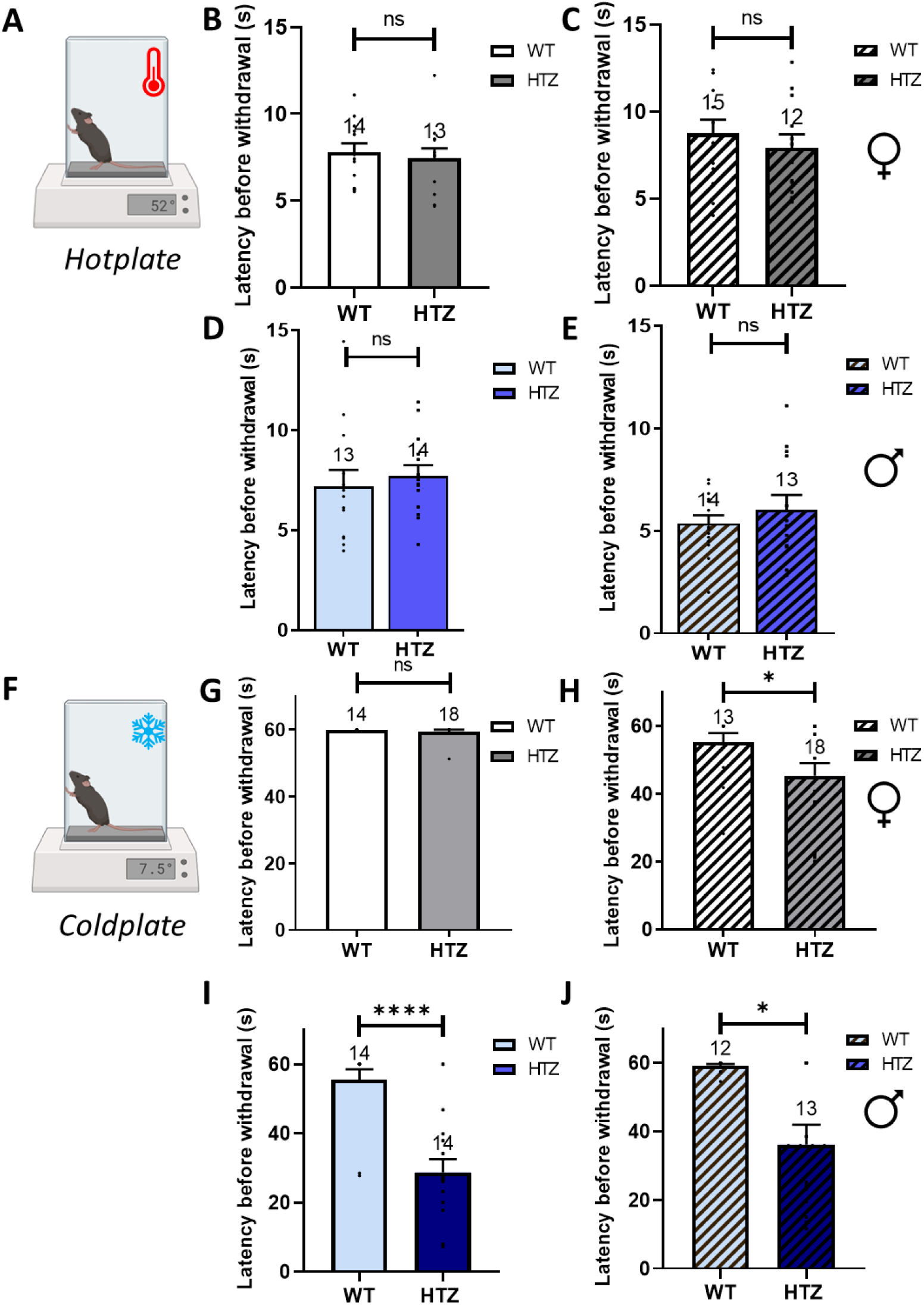
Evaluation of thermal sensitivity in juvenile and adult *Scn2a*^+/−^ (HTZ) mice. **A;F:** Schematic representation of the Hotplate (top, 52°c, 45s cut-off time) and Coldplate (bottom, 7.5°c, 1min cut-off time) test protocols. **B-E:** Heat sensitivity in female (B-C) and male (D-E) mice, left column: juveniles and right column: adults. **G-J**: Cold sensitivity in female (G-H) and male (I-J) mice, left column: juveniles and right column: adults. Statistical analysis: One sample Wilcoxon test p>0.9999 (juvenile female), and Mann-Whitney or Student’s t-test. *p<0.05 compared to WT, ****p<0.0001, ns: non-significant. The number of mice (N) is indicated above the bars. White or light blue: WT (female or male respectively), grey or dark blue: HTZ (female or male respectively), No pattern: Juvenile, Striped: Adult.

In the hotplate test, we observed no differences in the latency of reaction to the thermal stimulus between HTZ and WT littermates at both ages, neither in females (juvenile 7.5±0.6s HTZ; 7.8±0.5s WT; Student’s t-test, t(25)=0.4541, p=0.65, NS; adult 7.9±0.8s HTZ; 8.7±0.8s WT: Student’s t-test, t(25)=0.7302, p=0.47, NS, Figure5B & C) nor in males (juvenile 7.7±0.5s HTZ; 7.2±0.8s WT; Student’s t-test: t(25)=0.5433, p=0.59, NS; adult 6.0±0.7s HTZ; 5.3±0.4s WT; Student’s t-test: t(25)=0.8448, p=0.41, NS, Figure5D & E).

The cold plate test revealed a significant difference between genotypes for males at both ages and in adult females. In fact, adult HTZ females showed about 20% reduction in the latency to the first sign of discomfort compared to WT littermates (juvenile 59.5±0.5s HTZ, 60s WT: one-sample Wilcoxon test for HTZ because all WT reached the cut-off time, 60s, p>0.99, NS; adult 45.4±3.7s HTZ; 55.2±2.8s WT; Mann-Whitney, U=69, p=0.0364 Figure5G & H). Both juvenile and adult HTZ males showed around 50% reduction in the latency to the first sign of discomfort compared to their WT littermates (juvenile 28.7±3.8s HTZ; 55.4±3.1s WT; Mann-Whitney: U=16, p<0.0001; adult 36.2±5.7s HTZ; 59.1±0.5s WT; Mann-Whitney: U=37.5, p=0.014, Figure5I & J).

Thus, male and female *Scn2a^+/−^* mice had normal heat sensitivity at both juvenile and adult ages, whereas male *Scn2a^+/−^* mice at both ages and adult *Scn2a^+/−^* female mice show hyper-reactivity to cold stimuli.

### *Scn2a^+/−^* female mice display normal recognition memory but altered spatial working memory and decision-making

Cognitive defects are frequently present in ASD patients and mouse models, including memory deficits. We assessed recognition memory with the novel object recognition test (NORT) and spatial working memory with the Y maze test, as these tests often disclose cognitive dysfunctions in ASD mouse models (Figure 6A-D).

**Figure 6:**
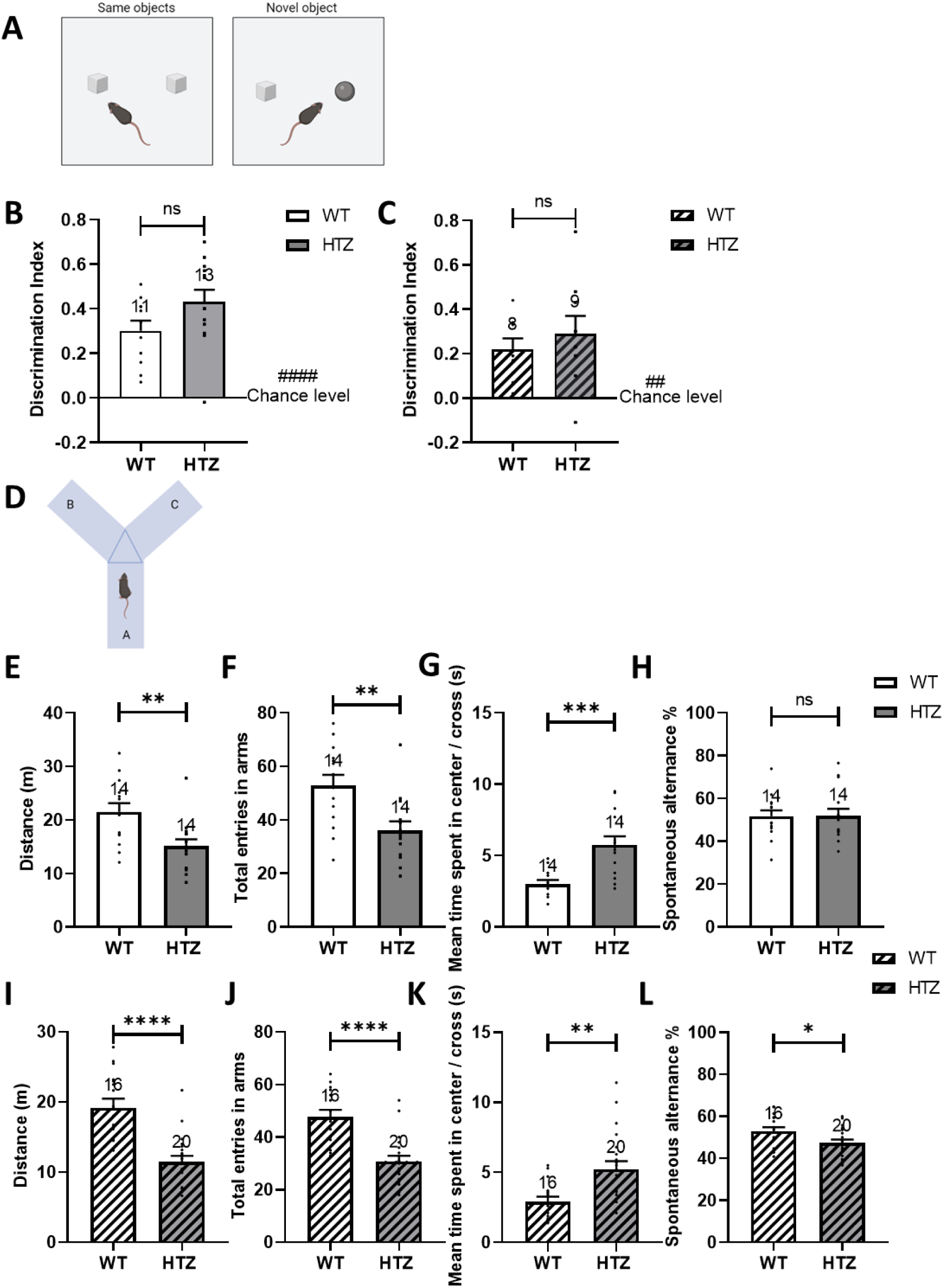
Evaluation of cognitive function in juvenile and adult female *Scn2a*^+/−^ (HTZ) mice. **A:** Schematic representation of the Novel Object Recognition Test (NORT) protocol divided in 2 phases: Learning phase with 2 identical objects and Object Recognition phase with a familiar and a new unfamiliar object. **B-C:** Discrimination index of the NORT for juvenile (B) and adult (C) female mice. **D:** Schematic representation of the Y maze apparatus. Quantifications for juvenile (E-H) and adult (I-L) female mice. **E-G:** Exploration parameters in the Y maze apparatus: Distance (E), total entries in the arms (F) and mean time spent in the center per cross (G) in juvenile female mice. **H:** Percent spontaneous alternation in juvenile female mice. **I-K:** Exploration parameters in the Y maze apparatus: Distance (I), total entries in the arms (J) and mean time spent in the center per cross (K) in adult female mice. **L:** Percent spontaneous alternation in adult female mice. Statistical analysis: Student’s t-test. *p<0.05 compared to WT, **p<0.01, ***p<0.001, ****p<0.0001, ns: non-significant. The number of mice (N) is indicated above the bars. White: WT, Grey: HTZ, No pattern: Juvenile, Striped: Adult.

We found no significant differences in the discrimination index of the NORT for both juvenile (0.43±0.05 HTZ; 0.30±0.05 WT; Student’s t-test: t(22)=1.825, p=0.825, NS, Figure 6B) and adult females (0.29±0.08 HTZ; 0.22±0.05 WT; Student’s t-test: t(15)=0.7086, p=0.4894, Figure 6C), which is consistent with normal recognition memory.

For the Y maze test, we first investigated the parameters that are usually used to quantify locomotor behavior in this test. Both juvenile and adult HTZ females showed a decrease in the total distance traveled (juvenile: 15.1±1.3m HTZ; 21.5±1.6m WT; t(26)=3.081, Student’s t-test: p=0.0048, Figure 6E; adult: 11.5±0.8m HTZ; 19.2±1.3m WT; Student’s t-test: t(34)=5.210, p<0.0001, Figure 6I) and a decrease in the total number of entries in the three arms of the maze (juvenile: 36.1±3.3 HTZ; 52.8±4.1 WT; Student’s t-test: t(26)=3.176, p=0.0038, Figure 6F; adult: 30.8±2.2 HTZ; 47.7±2.8 WT; Student’s t-test: t(34)=4.852, p<0.0001, Figure 6J). This result was initially surprising, because we did not find modifications of locomotor activity in other tests. To shed more light on this issue, we quantified the average time that mice spent in the center zone before entering a novel arm, which may be interpreted as a measure of the decision-making duration in the maze^45^. We observed about two-fold increase in the time spent in the center in HTZ females compared to WT littermates at both ages ((juvenile: 5.7±0.6s HTZ; 3.0±0.3s WT; Student’s t-test: t(26)=4.113, p=0.0003, Figure 6G; adult: 5.2±0.6s HTZ; 2.9±0.3s WT; Student’s t-test: t(34)=3.140, p=0.0035, Figure 6K). Finally, we identified in adult HTZ females, but not in juvenile ones, a little but significant reduction of the percentage of spontaneous alternations compared to WT littermates (juvenile: 51.9±3.3% HTZ; 51.7±2.8% WT; Student’s t-test: t(26)=0.04132, p=0.97, NS, Figure 6H; adult: 47.3±1.6% HTZ; 52.9±1.9% WT; Student’s t-test: t(34)=2.236, p=0.032, Figure 6L).

Hence, *Scn2a^+/−^*female mice have normal recognition memory at both ages, but show reduced exploratory behavior in a decision-making context, along with a slight reduction in performance in a spatial working memory task in adult females.

### Juvenile *Scn2a^+/−^* mice have reduced anxiety-like behavior in the Elevated Plus Maze test

Anxiety is a comorbidity often reported in ASD patients. We studied anxiety-like behavior in both juvenile and adult female mice in the Elevated Plus Maze (EPM) test: a cross-shaped maze elevated from the ground with two closed (safe) arms and two open (anxiogenic) arms. Anxiety-like behavior is determined evaluating the number of entries and fraction of time spent in the open arms (Figure7A). We initially evaluated global locomotion and exploration in the EPM by quantifying the total distance traveled, which was not different compared to control littermates (juvenile: 10.3±0.7m HTZ; 11.1±0.9m WT; Mann-Whitney test, U=115, p=0.69, NS, Figure 7C; adult: 9.3±0.5m HTZ; 9.3±0.5m WT; Student’s t-test: t(33)=0.0042, p=0.99, NS, Figure 7F). In juvenile females, we observed a significant increase in the percentage of entries in the open arms (54.3±2.2% HTZ; 44.1±2.4% WT; Student’s t-test, t(30)=3.080, p=0.0044, Figure 7D) and in the percentage of time spent in the open arms (53.1±3.1% HTZ; 35.1±3.1% WT; Student’s t-test, t(30)=4.055, p=0.0003, Figure 7G). In adult females, we observed strong trends towards an increase of the percentage of entries in the open arms (50.7±3.5% HTZ; 43.7±1.9% WT; Student’s t-test with Welch correction, t(23.81)=1.757, p=0.0918, Figure 7E) and in the percentage of time spent in the open arms (44.4±5.6% HTZ; 31.8±2.8% WT; Student’s t-test with Welch correction, t(22.36)=2.005, p=0.0572, Figure 7H). Thus, juvenile *Scn2a^+/−^*female mice showed reduced anxiety in the EPM, but normal locomotor and exploratory features. Adult *Scn2a^+/−^* female mice showed a reduction of the severity of this dysfunction, as we previously observed for male mice^23^.

**Figure 7:**
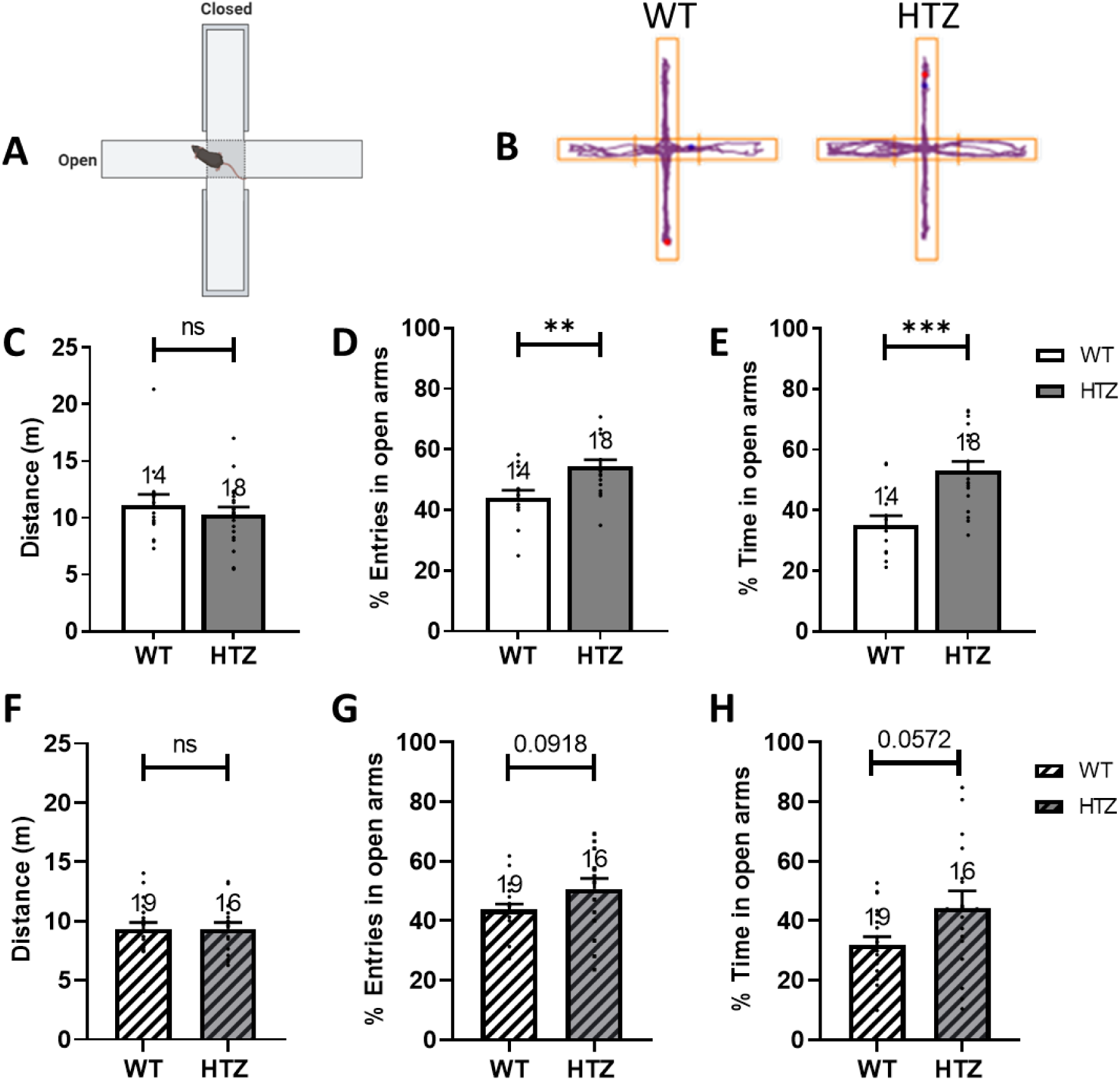
Evaluation of anxiety-like behavior in juvenile and adult *Scn2a*^+/−^ (HTZ) female mice. **A:** Schematic representation of the Elevated Plus-Maze (EPM) test. **B:** Representative trackplots of WT and HTZ female. **C-E:** Results in juvenile female mice: Distance in the maze (C), percent of entries (D) and time (E) spent in open arms of the EPM compared to total open + close arms entries or time. **F-H:** Results in adult female mice: Distance in the maze (F), percent of entries (G) or time (H) spent in open arms of the EPM compared to total open + close arms entries or time. Statistical analysis: Mann-Whitney or Student’s t-test (with Welch’s correction for G-H). *p<0.05 compared to WT, **p<0.01, ***p<0.001, ns: non-significant. The number of mice (N) is indicated above the bars. White: WT, Grey: HTZ, Empty: Juvenile and Striped: Adult.

## Discussion

We performed an extended characterization of the behavioral dysfunctions in female *Scn2a^+/−^* mice, for which little information is available. Because ASD is a neurodevelopmental disorder with onset in childhood, we investigated both juvenile (P21-42) and adult mice (>P70), as we have already done for male mice^23^. Some recent studies of mouse models of *SCN2A* genetic variants have included female mice in their investigations^27,29^, but they have not performed a global behavioral assessment and they analyzed only late developmental stages.

Dysfunction of social behaviors is among the core symptoms of ASD. We observed that *Scn2a^+/−^* female mice display an increased sociability in the 3-chamber test both at juvenile and adult ages, without alterations of social memory and without modifications of olfactory discrimination of social or non-social cues. Increased sociability in the 3-chamber test and in direct social interaction was previously reported in *Scn2a^+/−^*adult male mice^26,30^, as well as in adult male mice of other models of *SCN2A* haploinsufficiency and ASD-like phenotypes^28,29^. Interestingly, it has been proposed that the hyper-social phenotype observed in *Scn2a^Δ1898/+^* mouse model could model the inappropriate social approach behavior reported in the Simons Foundation database for ASD patients with *SCN2A* variants^29^. In our previous study of *Scn2a^+/−^* male mice, we have observed decreased social communication by ultra-sonic vocalizations (USVs) at both young and adult age^23^, which is considered an autistic-like trait. We have not evaluated USVs in *Scn2a^+/−^* female mice, because females generate much less USVs than males^46–48^, complicating the quantification. In particular, in a male-female encounter context, which we have used for adult *Scn2a*^+/−^ male mice^23^, USVs are produced by males^49,50^.

Restricted, stereotyped and repetitive behaviors are other core symptoms of ASD. We assessed the presence of innate motor stereotypies in *Scn2a^+/−^* female mice by measuring the amount of time spent self-grooming in an open field arena and the number of marbles buried, which were increased in *Scn2a^+/−^* males^23,51^. Instead, in *Scn2a^+/−^* female mice we have not observed any differences in these tests compared to WT. Moreover, we found that *Scn2a^+/−^* female mice do not show a more pronounced acquisition of repetitive motor routines evaluated with the rotarod test, feature that we instead observed in *Scn2a^+/−^* male mice^23^. Interestingly, this gender difference was observed also in the *Scn2a^K1422E^*model of *Scn2a* haploinsufficiency and ASD-like phenotype^28^. Overall, our results do not show acquired or innate motor stereotypies in *Scn2a^+/−^*female mice. Moreover, we did not observe alterations of exploratory and locomotor behavior in the open field test, indicating that *Scn2a^+/−^* female mice do not show general hyperactivity, which is a frequent comorbidity of ASD patients and mouse models.

Abnormalities in sensory processing are further features of ASD phenotypes. Because in our previous study of male *Scn2a^+/−^* mice we have not investigated these features^23^, we have here directly compared the response to thermal stimuli in male and female *Scn2a^+/−^* mice. Both juvenile and adult male *Scn2a^+/−^*mice showed hyper-reactivity to cold stimuli, whereas female *Scn2a^+/−^*mice showed hyper-reactivity only in adulthood, and neither males nor females showed altered response to heat. This is consistent with a milder phenotype in females. Notably, a recent study showed hyper-reactivity to both cold and hot stimuli in gene-trap *Scn2a^gtKO/gtKO^* mice^27^, which may be caused by the larger reduction of *Scn2a* expression in that model compared to haploinsufficient *Scn2a^+/−^* mice.

Increased anxiety is a frequent comorbidity in ASD patients and mouse models. We observed instead an apparent decrease in anxiety-like phenotype: more entries and more time spent in the open anxiogenic areas of the elevated plus maze apparatus in juvenile *Scn2a^+/−^* female mice and a trend towards an increase in adults. Interestingly, these results are consistent with the ones previously observed in *Scn2a^+/−^* male^23,26^ and female mice^31^. Moreover, decreased anxiety has also been identified in other ASD mouse models, such as *Scn2a^K1422E^* ^28^, *Shank3b*^−/−^, *Caspr2*^−/−^ and 16p11.2 mice^52^. However, this may be related to a dysfunctional risk-assessment behavior, which would lead to comparable exploration of risky and non-risky areas, as already proposed for other ASD models^52^. Notably, decreased risk-assessment and impulsivity have been observed also in ASD patients, leading in some cases to life threatening behaviors^53–56^.

Cognitive functions are often impaired in ASD patients and mouse models. Similar to social memory discussed above, neither juvenile nor adult *Scn2a^+/−^*female mice showed dysfunctions of recognition memory in the NORT. Thus, unlike in males, we found an intact learning and memory storage in *Scn2a^+/−^*female mice, in both short (10min in the 3-chambers test) and long (1h in the NORT) periods of time. However, in the Y maze test, we found a mild decrease of spatial working memory (spontaneous alternations) in adult females, which seems gender-specific, since *Scn2a*^+/−^ males show a normal spatial working memory both at young^23^ and adult^23,31^ age. In this test, we also observed an impairment in exploratory parameters (distance travelled and total arms entries) in both juvenile and adult females. The difference seen in spontaneous alternations cannot be explained by a reduction in exploratory behavior, because there is no correlation between the number of arm entries and the percentage of spontaneous alternations^57^. Notably, the decreased distance travelled and exploration in the Y maze are not caused by a locomotor deficit, which was not observed in other tests. Evaluating this issue, we found in *Scn2a*^+/−^ females a 2-fold increase in the time spent in the center of the maze at both ages, which resembled to a “hesitation” of the animal to choose where to go. We interpreted this behavioral abnormality as altered decision-making. In fact, it has been proposed that, in maze tasks involving arm choices, rodents show pauses in choice-related points that are related to the decision-making process, called vicarious trial-and-error^45^. The observed behavior of *Scn2a^+/−^* female mice in the Y-maze test was consistent with a longer decision-making time, which was followed by a relatively high percentage of correct choices in alternation but led to a decrease in the distance travelled and in the total arms entries. This feature is consistent with the atypical decision-making of ASD patients in choice tests^58^. Therefore, we observed in *Scn2a^+/−^* female mice a reduction in anxiety-like behaviors in juveniles, hypersensitivity to cold stimuli and a slight reduction in spatial working memory in adulthood, as well as social behavior abnormalities and decision-making deficits at both ages (Table).

**Table:**
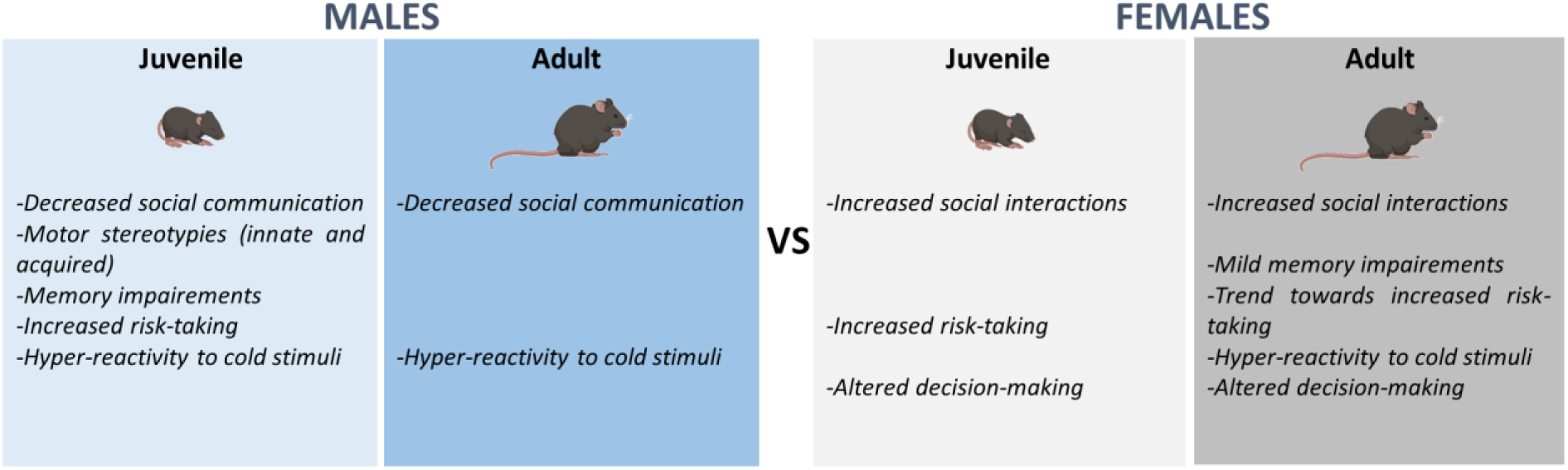
Comparison of behavioral abnormalities between *Scn2a^+/−^* male and female mice at juvenile and adult age that we observed in our studies (this report and ^23^)

| MALES |  | FEMALES |  |
| --- | --- | --- | --- |
| Juvenile | Adult | Juvenile | Adult |
| -Decreased social communication<br>-Motor stereotypies (innate and acquired)<br>-Memory impairments<br>-Increased risk-taking<br>-Hyper-reactivity to cold stimuli | -Decreased social communication<br>-Hyper-reactivity to cold stimuli | -Increased social interactions<br>-Increased risk-taking<br>-Altered decision-making | -Increased social interactions<br>-Mild memory impairments<br>-Trend towards increased risk-taking<br>-Hyper-reactivity to cold stimuli<br>-Altered decision-making |

In our previous study^23^ (Table), we observed in *Scn2a^+/−^* male mice a prominent ASD-like phenotype in juveniles, comprising social communication deficits (but without interaction or olfactory discrimination deficits), innate (increased marbles buried and time spent self-grooming) and acquired (increased exercise-induced latency to fall from the rotarod) stereotyped behavior, decreased anxiety-like and depressive-like behaviors associated to normal locomotion, functional spatial working and long-term memories but decreased object recognition memory. In adult *Scn2a*^+/−^ male mice, the only behavioral abnormality we identified was a decreased social communication.

Other studies have investigated behavioral features of *Scn2a* models. A study reported in *Scn2a*^+/−^ adult male mice a delayed learning in spatial working memory tasks^24^, which could be related to the working memory defect we observed in adult females in the Y-maze test. Another study reported in adult *Scn2a^+/−^*male mice novelty-induced hyperactivity in the openfield, decreased social approach in the resident-intruder test, increased dominance in the tube test, conflicting results in depressive-like behavior in the force swim test and the tail suspension test, enhanced fear learning and decreased extinction^26^. A further study reported very mild phenotypes in adult *Scn2a*^+/−^ mice, both males and females, with mild non-significant trends toward deficits in learning and social behavior and significant effects only for decreased social memory and increased risk taking in females^31^. A study of another *Scn2a* knock-out model (deletion of exons 4-6) identified in adult male mice altered spatial learning and fear extinction abnormalities associated to enhanced direct social interaction but no social communication or motor stereotypies^30^. In hypomorphic *Scn2a^Δ1898/+^* adult mice it has been recently reported in adult males but not females an increased sociability in the 3-chambers test^29^, which resembles the increased sociability that we observed here in female *Scn2a^+/−^*mice. Thus, there is some inter-laboratory variability in the results of behavioral studies, which may be related to specific breeding, handling and experimental design conditions and that complicates the comparison of detailed features. However, a general finding is that adult *Scn2a* loss-of-function models have relatively mild behavioral phenotypes.

Overall, our results are consistent with the literature, showing that both male and female adult *Scn2a*^+/−^ mice have mild phenotypes. Additionally, we have shown that, differently than *Scn2a*^+/−^ male mice, also in the juvenile period *Scn2a*^+/−^ female mice have a milder phenotype, whose features are therefore less markedly reduced in adulthood. Moreover, we have observed some gender-specific features. Interestingly, a milder phenotype of females compared to males has also been observed in other ASD mouse models^59,60^.

As already outlined in the introduction, it has been shown that ASD female patients can express symptoms differently than men and have more prominent masking behaviors^9,12–15^, leading to a delayed or incorrect diagnosis and an underestimation of ASD cases in ^9–11^. Our data obtained in *Scn2a*^+/−^ mice are consistent with biological mechanisms for the specific features of ASD in females.

There is little information in the literature about the detailed phenotype of ASD patients carrying *SCN2A* variants, in particular regarding gender-related features. The few female cases described in some details presented with developmental and epileptic encephalopathies or other neurodevelopmental disorders^61–64^, consistent with a more difficult identification of female ASD phenotypes. Notably, although overall mild, we have found some robust behavioral features in *Scn2a^+/−^* mice.

In conclusion, our study identifies for the first time the global behavioral phenotype of juvenile and adult *Scn2a^+/−^* female mice and compares it with that of *Scn2a^+/−^* male mice. The specific features and the relatively mild phenotype of *Scn2a^+/−^* female mice highlight the importance of including females in the study of animal models and developing gender-specific clinical assessments for ASD patients.

## Acknowledgements.

We thank Emilie Bonnet (Institute of Molecular and Cellular Pharmacology, CNRS UMR 7275 and University Cote d’Azur) for her skillful technical support on colony management/genotyping, Eric Lingueglia/Emmanuel Deval’s team (Institute of Molecular and Cellular Pharmacology, CNRS UMR 7275 and University Cote d’Azur) for sharing some behavioral apparatus and Edward Glasscock for providing founders for the colony of *Scn2a^+/−^*mice.

## Author Contributions

I.L. and M.M. conceived the project. W.M. and M.S. performed the experiments and analyzed the data. W.M. wrote an initial draft of the paper. M.S, I.L. and M.M critically revised the manuscript. All authors approved the final version of the manuscript.

## Data Availability Statement

Data are available on request.

## References

1. American Psychiatric Association. Diagnostic and statistical manual of mental disorders. 5th ed.

2. Rylaarsdam, L. & Guemez-Gamboa, A. Genetic Causes and Modifiers of Autism Spectrum Disorder. Front. Cell. Neurosci. 13, (2019).

3. Lai, M.-C., Lombardo, M. V. & Baron-Cohen, S. Autism. Lancet Lond. Engl. 383, 896–910 (2014).

4. Maenner, M. J. Prevalence and Characteristics of Autism Spectrum Disorder Among Children Aged 8 Years — Autism and Developmental Disabilities Monitoring Network, 11 Sites, United States, 2018. MMWR Surveill. Summ. 70, (2021).

5. Fombonne, E. Epidemiological surveys of autism and other pervasive developmental disorders: an update. J. Autism Dev. Disord. 33, 365–382 (2003).

6. Loomes, R., Hull, L. & Mandy, W. P. L. What Is the Male-to-Female Ratio in Autism Spectrum Disorder? A Systematic Review and Meta-Analysis. J. Am. Acad. Child Adolesc. Psychiatry 56, 466–474 (2017).

7. Kirkovski, M., Enticott, P. G. & Fitzgerald, P. B. A review of the role of female gender in autism spectrum disorders. J. Autism Dev. Disord. 43, 2584–2603 (2013).

8. Van Wijngaarden-Cremers, P. J. M. et al. Gender and age differences in the core triad of impairments in autism spectrum disorders: a systematic review and meta-analysis. J. Autism Dev. Disord. 44, 627–635 (2014).

9. Hull, L. et al. “Putting on My Best Normal”: Social Camouflaging in Adults with Autism Spectrum Conditions. J. Autism Dev. Disord. 47, 2519–2534 (2017).

10. Kočovská, E. et al. The Rising Prevalence of Autism: A Prospective Longitudinal Study in the Faroe Islands. J. Autism Dev. Disord. 42, 1959–1966 (2012).

11. Lai, M.-C. & Szatmari, P. Sex and gender impacts on the behavioural presentation and recognition of autism. Curr. Opin. Psychiatry 33, 117–123 (2020).

12. Gould, J. & Ashton-Smith, J. Missed diagnosis or misdiagnosis? Girls and women on the autism spectrum. Good Autism Pract. GAP 12, 34–41 (2011).

13. Lai, M.-C. et al. Quantifying and exploring camouflaging in men and women with autism. Autism Int. J. Res. Pract. 21, 690–702 (2017).

14. McQuaid, G. A., Lee, N. R. & Wallace, G. L. Camouflaging in autism spectrum disorder: Examining the roles of sex, gender identity, and diagnostic timing. Autism Int. J. Res. Pract. 26, 552–559 (2022).

15. Rynkiewicz, A. et al. An investigation of the ‘female camouflage effect’ in autism using a computerized ADOS-2 and a test of sex/gender differences. Mol. Autism 7, 10 (2016).

16. Lord, C. et al. Autism spectrum disorder. Nat. Rev. Dis. Primer 6, 1–23 (2020).

17. Sohal, V. S. & Rubenstein, J. L. R. Excitation-inhibition balance as a framework for investigating mechanisms in neuropsychiatric disorders. Mol. Psychiatry 24, 1248–1257 (2019).

18. Mantegazza, M., Cestèle, S. & Catterall, W. A. Sodium channelopathies of skeletal muscle and brain. Physiol. Rev. 101, 1633–1689 (2021).

19. Rusina, E., Simonti, M., Duprat, F., Cestèle, S. & Mantegazza, M. Voltage-gated sodium channels in genetic epilepsy: up and down of excitability. J. Neurochem. n/a,.

20. Sanders, S. J. et al. Progress in Understanding and Treating SCN2A-Mediated Disorders. Trends Neurosci. 41, 442–456 (2018).

21. Ben-Shalom, R. et al. Opposing Effects on NaV1.2 Function Underlie Differences Between SCN2A Variants Observed in Individuals With Autism Spectrum Disorder or Infantile Seizures. Biol. Psychiatry 82, 224–232 (2017).

22. Planells-Cases, R. et al. Neuronal death and perinatal lethality in voltage-gated sodium channel alpha(II)-deficient mice. Biophys. J. 78, 2878–2891 (2000).

23. Léna, I. & Mantegazza, M. NaV1.2 haploinsufficiency in Scn2a knock-out mice causes an autistic-like phenotype attenuated with age. Sci. Rep. 9, 12886 (2019).

24. Middleton, S. J. et al. Altered hippocampal replay is associated with memory impairment in mice heterozygous for the Scn2a gene. Nat. Neurosci. 21, 996–1003 (2018).

25. Ogiwara, I. et al. Nav1.2 haplodeficiency in excitatory neurons causes absence-like seizures in mice. Commun. Biol. 1, (2018).

26. Tatsukawa, T. et al. Scn2a haploinsufficient mice display a spectrum of phenotypes affecting anxiety, sociability, memory flexibility and ampakine CX516 rescues their hyperactivity. Mol. Autism 10, 15 (2019).

27. Eaton, M. et al. Generation and basic characterization of a gene-trap knockout mouse model of Scn2a with a substantial reduction of voltage-gated sodium channel Nav 1.2 expression. Genes Brain Behav. 20, e12725 (2021).

28. Echevarria-Cooper, D. M. et al. Cellular and behavioral effects of altered NaV1.2 sodium channel ion permeability in Scn2aK1422E mice. Hum. Mol. Genet. ddac087 (2022) doi:10.1093/hmg/ddac087.

29. Wang, H.-G. et al. Scn2a severe hypomorphic mutation decreases excitatory synaptic input and causes autism-associated behaviors. JCI Insight 6, 150698 (2021).

30. Shin, W. et al. Scn2a Haploinsufficiency in Mice Suppresses Hippocampal Neuronal Excitability, Excitatory Synaptic Drive, and Long-Term Potentiation, and Spatial Learning and Memory. Front. Mol. Neurosci. 12, 145 (2019).

31. Spratt, P. W. E. et al. The Autism-Associated Gene Scn2a Contributes to Dendritic Excitability and Synaptic Function in the Prefrontal Cortex. Neuron 103, 673–685.e5 (2019).

32. Lee, T. T.-Y. & Gorzalka, B. B. Timing is everything: evidence for a role of corticolimbic endocannabinoids in modulating hypothalamic-pituitary-adrenal axis activity across developmental periods. Neuroscience 204, 17–30 (2012).

33. Brust, V., Schindler, P. M. & Lewejohann, L. Lifetime development of behavioural phenotype in the house mouse (Mus musculus). Front. Zool. 12, S17 (2015).

34. Crawley, J. N. Designing mouse behavioral tasks relevant to autistic-like behaviors. Ment. Retard. Dev. Disabil. Res. Rev. 10, 248–258 (2004).

35. Moy, S. S. et al. Sociability and preference for social novelty in five inbred strains: an approach to assess autistic-like behavior in mice. Genes Brain Behav. 3, 287–302 (2004).

36. Witt, R. M., Galligan, M. M., Despinoy, J. R. & Segal, R. Olfactory behavioral testing in the adult mouse. J. Vis. Exp. JoVE 949 (2009) doi:10.3791/949.

37. Brielmaier, J. et al. Autism-relevant social abnormalities and cognitive deficits in engrailed-2 knockout mice. PloS One 7, e40914 (2012).

38. Thomas, A. et al. Marble burying reflects a repetitive and perseverative behavior more than novelty-induced anxiety. Psychopharmacology (Berl.) 204, 361–373 (2009).

39. Antunes, M. & Biala, G. The novel object recognition memory: neurobiology, test procedure, and its modifications. Cogn. Process. 13, 93–110 (2012).

40. Hamadate, N. et al. Liposome-Encapsulated Hemoglobin Ameliorates Impairment of Fear Memory and Hippocampal Dysfunction After Cerebral Ischemia in Rats. J. Pharmacol. Sci. 114, 409–419 (2010).

41. Ohmura, Y. et al. Disruption of model-based decision making by silencing of serotonin neurons in the dorsal raphe nucleus. Curr. Biol. 31, 2446–2454.e5 (2021).

42. Nunes. Axonal sodium channel NaV1.2 drives granule cell dendritic GABA release and rapid odor discrimination. PLoS biology (2018).

43. Rothwell, P. E. et al. Autism-associated neuroligin-3 mutations commonly impair striatal circuits to boost repetitive behaviors. Cell 158, 198–212 (2014).

44. Corder, G. et al. Loss of μ opioid receptor signaling in nociceptors, but not microglia, abrogates morphine tolerance without disrupting analgesia. Nat. Med. 23, 164–173 (2017).

45. Redish, A. D. Vicarious trial and error. Nat. Rev. Neurosci. 17, 147–159 (2016).

46. Sewell), G. D. S. (née. Ultrasound and mating behaviour in rodents with some observations on other behavioural situations. J. Zool. 168, 149–164 (1972).

47. White, N. R., Prasad, M., Barfield, R. J. & Nyby, J. G. 40- and 70-kHz vocalizations of mice (Mus musculus) during copulation. Physiol. Behav. 63, 467–473 (1998).

48. Whitney, G., Coble, J. R., Stockton, M. D. & Tilson, E. F. Ultrasonic emissions: do they facilitate courtship of mice. J. Comp. Physiol. Psychol. 84, 445–452 (1973).

49. Hanson, J. L. & Hurley, L. M. Female Presence and Estrous State Influence Mouse Ultrasonic Courtship Vocalizations. PLOS ONE 7, e40782 (2012).

50. Peleh, T., Eltokhi, A. & Pitzer, C. Longitudinal analysis of ultrasonic vocalizations in mice from infancy to adolescence: Insights into the vocal repertoire of three wild-type strains in two different social contexts. PloS One 14, e0220238 (2019).

51. Indumathy, J., Pruitt, A., Gautier, N. M., Crane, K. & Glasscock, E. Kv1.1 deficiency alters repetitive and social behaviors in mice and rescues autistic-like behaviors due to Scn2a haploinsufficiency. Brain Behav. 11, e02041 (2021).

52. Oron, O. et al. Gene network analysis reveals a role for striatal glutamatergic receptors in dysregulated risk-assessment behavior of autism mouse models. Transl. Psychiatry 9, 257 (2019).

53. Bornovalova, M. A. et al. Risk taking differences on a behavioral task as a function of potential reward/loss magnitude and individual differences in impulsivity and sensation seeking. Pharmacol. Biochem. Behav. 93, 258–262 (2009).

54. Guan, J. & Li, G. Injury Mortality in Individuals With Autism. Am. J. Public Health 107, 791–793 (2017).

55. Moeller, F. G., Barratt, E. S., Dougherty, D. M., Schmitz, J. M. & Swann, A. C. Psychiatric aspects of impulsivity. Am. J. Psychiatry 158, 1783–1793 (2001).

56. South, M., Dana, J., White, S. E. & Crowley, M. J. Failure is not an option: Risk-taking is moderated by anxiety and also by cognitive ability in children and adolescents diagnosed with an autism spectrum disorder. J. Autism Dev. Disord. 41, 55–65 (2011).

57. Miedel, C. J., Patton, J. M., Miedel, A. N., Miedel, E. S. & Levenson, J. M. Assessment of Spontaneous Alternation, Novel Object Recognition and Limb Clasping in Transgenic Mouse Models of Amyloid-β and Tau Neuropathology. JoVE J. Vis. Exp. e55523 (2017) doi:10.3791/55523.

58. Mosner, M. G. et al. Vicarious Effort-Based Decision-Making in Autism Spectrum Disorders. J. Autism Dev. Disord. 47, 2992–3006 (2017).

59. Jeon, S. J. et al. Sex-specific Behavioral Features of Rodent Models of Autism Spectrum Disorder. Exp. Neurobiol. 27, 321–343 (2018).

60. Schaafsma, S. M. et al. Sex-specific gene–environment interactions underlying ASD-like behaviors. Proc. Natl. Acad. Sci. 114, 1383–1388 (2017).

61. D’Gama, A. M. et al. Targeted DNA Sequencing from Autism Spectrum Disorder Brains Implicates Multiple Genetic Mechanisms. Neuron 88, 910–917 (2015).

62. Fromer, M. et al. De novo mutations in schizophrenia implicate synaptic networks. Nature 506, 179–184 (2014).

63. Suddaby, J. S., Silver, J. & So, J. Understanding the schizophrenia phenotype in the first patient with the full SCN2A phenotypic spectrum. Psychiatr. Genet. (2019) doi:10.1097/YPG.0000000000000219.

64. Yamamoto, T. et al. Genomic backgrounds of Japanese patients with undiagnosed neurodevelopmental disorders. Brain Dev. 41, 776–782 (2019).

